# A High-Resolution LED Display for Steady State Visual Stimulation: Customized, Affordable, and Open Source

**DOI:** 10.1101/2023.12.04.569998

**Authors:** Monica Otero, Yunier Prieur-Coloma, Wael El-Deredy, Alejandro Weinstein

## Abstract

Visually evoked steady-state potentials (SSVEPs) are neural responses elicited by visual stimuli oscillating at specific frequencies. In this study, we introduce a novel LED display system designed specifically for steady-state visual stimulation, offering precise control over visual stimulus parameters, including frequency resolution, luminance, and the ability to control the phase at the end of the stimulation. The LED display provides a personalized, modular, and affordable option for experimental setups. Based on the Teensy 3.2 board, the display utilizes Direct Digital Synthesis and Pulse Width Modulation techniques to control the LEDs. Its performance is validated through four experiments: the first two measure LED light intensities directly, while the last two assess the display’s impact on EEG recordings. The results demonstrate that the display can deliver a stimulus suitable for generating SSVEPs with the desired frequency and phase resolution. We provide comprehensive documentation, including all necessary codes and electrical diagrams, as an open-source resource. This facilitates the replication and adaptation of the system for specific experimental requirements, enhancing its potential for widespread use in the field of neuroscience.

## 1. Introduction

Sensory repetitive stimulation induces neural activity synchronization in populations of neurons of the cortex, which is reflected as distinct oscillatory patterns in the electroencephalogram (EEG) across various sensory modalities. In the case of visual stimulation, Steady State Visual Evoked Potentials (SSVEPs) are neural responses elicited by visual stimuli oscillating at specific frequencies, providing valuable insights into the functional organization of visual processing and cognitive functions in health [1–4] and pathological conditions [5–10]. SSVEPs can be recorded from the scalp using the EEG, obtaining maximum amplitude at the stimulation frequency, usually at occipital brain regions. The amplitude of frequency components in SSVEP signals remains consistent over time, allowing for reliable identification of stimulus frequency through frequency domain analysis [11]. The robust frequency characteristics of SSVEPs have led to the widespread utilization of the frequency tagging technique, which involves encoding multiple visual targets with distinct flickering frequencies [12,13]. This technique has found extensive applications in visual neuroscience and neural engineering domains.

To investigate SSVEPs, precise and customizable visual stimulation is essential. However, conventional displays have fixed sampling rates, limiting the number of frequencies that can be presented and potentially missing critical frequency components. The drawbacks mentioned above and the high cost of acquiring customizable Light-Emitting Diodes (LED) displays present significant challenges in experimental setups, limiting accessibility for numerous researchers.

Different types of visual stimuli can generate SSVEPs, including flickering checker-board patterns, sinusoidal gratings, or even complex images modulated at specific frequencies [1]. Each type of stimulus has unique advantages and applications, enabling researchers to investigate various aspects of visual processing and cognitive functions. This article introduces a customizable and affordable LED display to generate sinusoidal-like visual stimuli for SSVEPs. This display is designed to provide a high degree of control over the parameters of the visual stimuli, including their luminance, frequency resolution, and wave phase. It also offers a range of customization options, allowing researchers to tailor the stimuli to their specific experimental needs.

Pulse Width Modulation (PWM) is a modulation technique that generates variable-width pulses to encode the amplitude of an analog input signal [14]. Direct Digital Synthesis (DDS) is a technique to generate arbitrary analog signals using a digital circuit [15]. Combining these techniques, the LED display can produce visual stimulation signals resembling sinusoidal waves with precise high-frequency resolution and end phase, with the advantage of requiring a simple LED driving circuit.

Controlling the phase of the visual stimulation is crucial for various experimental paradigms, particularly in electrophysiological studies. The ability to control the phase of the stimulation allows for the investigation of phase-dependent neural responses and their relationship to cognitive processes [16–20].

In this article, leveraging the DDS and PWM techniques, we successfully produced visual stimuli at different frequencies while controlling the phase of both the initiation and termination of sinusoidal-like signals emitted by the LEDs. Additionally, the LED display described in this study enables the synchronization with external EEG equipment, allowing triggers to be recorded alongside neural activity. This feature is essential for accurate data analysis.

Furthermore, this article discusses the limitations of traditional displays, presenting a comprehensive exploration of the LED display’s technical properties, including both hardware and software components. Additionally, the advantages of employing a customizable LED display are highlighted, emphasizing the significance of phase control and synchronization with external EEG recording equipment in advancing SSVEP research. It is noteworthy that source codes and hardware schematics are made available as an open-source LED display design for periodic stimulation.

Due to its ease of use, flexibility, and potential for producing more reliable and consistent results, we believe this new tool can advance our understanding of visual processing and contribute to developing new approaches for diagnosing and researching several neurological disorders.

## 2. Materials and Methods

### 2.1. LED Display

The LED display has been designed to facilitate the development of entrainment experiments by presenting a sinusoidally varying light at a specific frequency customized to each participant and terminating at different phases of the sinusoid. The proposed display consists of two major components: a display controller and an array of four entrainer LEDs, as shown in Figure 1-(a). The display controller was designed using a Teensy 3.2 USB-based microcontroller development system (PJRC, OR, USA) that drives the entrainer LEDs using a PWM digital output and a transistor. In addition, a set of digital inputs and outputs is available to suit the needs of specific experiments. The array of entrainer LEDs generates visual stimulus for the experiments. It consists of four white LEDs, part number YSL-R1042WC-D15(China Yunsun LED Lighting Co., China), situated at the center of a 50×50 cm black display as vertices of a 5×5 cm square. The design can be modified to accommodate other LED arrangements or LED numbers. Additionally, the maximum intensity of the light emitted by the LEDs can be adjusted using a potentiometer.

**Figure 1.**
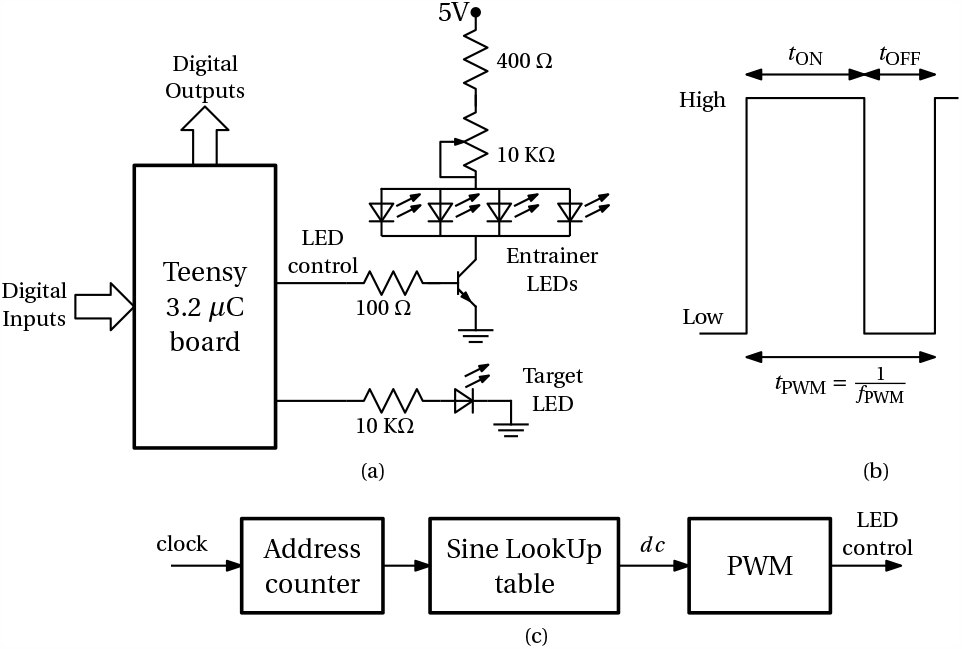
Block diagram of the display. (a) The Teensy 3.2 *μ*C board controls the display. The LED control signal is a PWM digital output that turns the entrainer LEDs on and off through a transistor. The LEDs maximum intensities are adjusted using a potentiometer. A digital output controls the target LED. A set of general digital input and output signals is also available. (b) The PWM signal has two levels: low (LEDs off) and high (LEDs on). The duty cycle is defined as 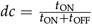. The PWM period is *t*_PWM_ = *t*_ON_ + *t*_OFF_, and its frequency is 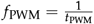. (c) The LED control signal is generated by combining DDS and PWM techniques. A clock signal increments an address counter. This counter is used to index a lookup table that encodes duty cycles with the values of each sine sample. The duty cycles are transformed into the digital signal that controls the LED by the PWM microcontroller peripheral.

The Teensy microcontroller and entrainer LEDs are powered by an external 5 V power supply.

We generated the LED control signal by combining the DDS [15] with the PWM [14] techniques (Fig. 1-(c)). The LED control signal is a PWM signal with a high-low pattern where a high level (LEDs are on) is followed by a low level (LEDs are off). In this pattern, the sum of the time the signal is high (*t*_ON_) with the time the signal is low (*t*_OFF_) is fixed, and it is called the PWM period *t*_PWM_, and its reciprocal is called the PWM frequency *f*_PWM_, as shown in Fig. 1-(b). For each PWM period, the duty cycle is defined as 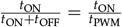. The duty cycle encodes the analog value to be generated. By having a high PWM frequency with respect to the visual system frequency response [21], this becomes a digital-to-analog conversion process. The duty cycle is generated using DDS, where the sinusoidal waveform with the desired frequency and end phase is stored in a lookup table. The lookup table values are read sequentially using an address counter incremented by one with each PWM period. The experiment stops when the address counter reaches the last value of the lookup table, or it is reset if one wishes to deliver the stimulus again.

We implemented the generation of the LED control signal in the Teensy 3.2 board using the functions `analogWriteFrequency` and `analogWrite` and the class `IntervalTimer` [22]. Functions `analogWriteFrequency` and `analogWrite` control the generation of the PWM signal. Class `IntervalTimer` creates a function that is called periodically. Listing 1 shows a code snippet with key implementation details (see supplementary material for the complete source code). The first line includes a header file with an array containing the sine lookup table values. Line 2 creates an `IntervalTimer dds` object that is used in line 6 to attach function `dds_update` to a timer, such that this function is executed every 1 ms. Line 5 uses `analogWriteFrequency` to set the PWM period *t*_PWM_ to 1 ms, or equivalently, to set the PWM frequency *f*_PWM_ to 1 kHz. Line 10 calls, from function `dds`, the function `analogWrite`.

This function sets the PWM duty cycle to the value `lookup_table[address_counter]`. Variable `ledPin` defines the Teensy pin used to generate the LED control signal. Lines 11 to 13 increment `address_counter` and stop the experiment if the counter reaches the last address (constant `MAX_ADDRESS`) of the lookup table (the experimenter can change line 13 to handle what to do after presenting the stimulus according to the experiment details).

#### Listing 1 Code snippet to generate the LED control signal

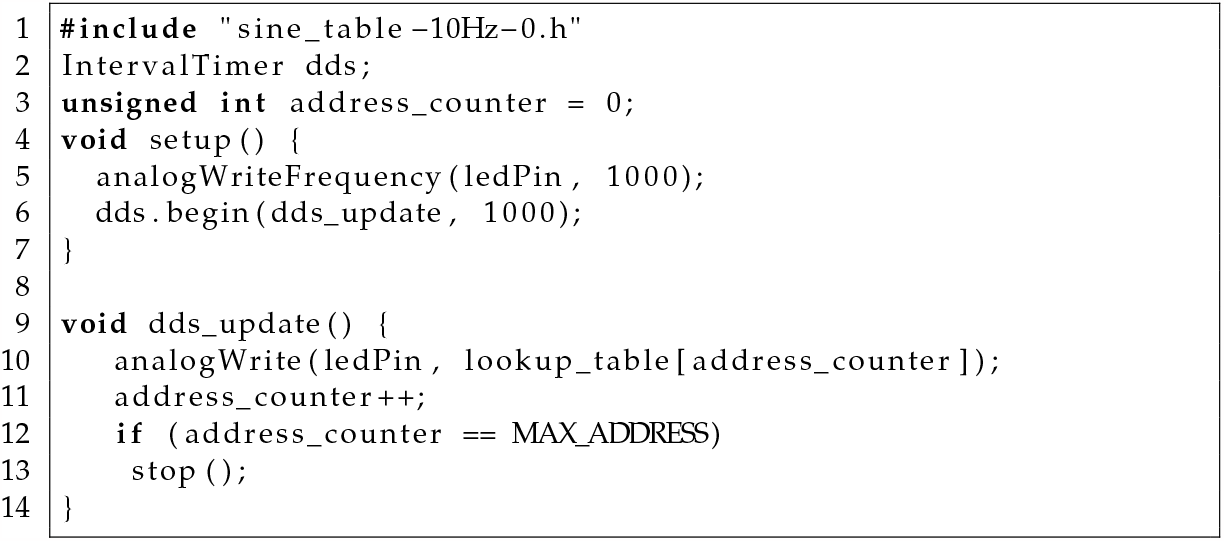

The function `analogWrite` sets the PWM with an integer value in the range of 0 to 256, where 0 corresponds to a duty cycle of 0, and 256 to a duty cycle of 1. For this reason, we fill the DDS lookup table with duty cycle (*dc*) values defined according to the following expression:

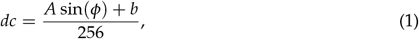

with *A* = 100 and *b* = 110, corresponding to duty cycles in the range 0.04 to 0.82, to avoid too narrow pulses. Variable *ϕ* is set according to the desired frequency and end phase. We provide a Python script (see Supplementary Materials) to create a header file with the lookup table corresponding to the desired stimulation parameters (frequency, end phase, and stimulation duration).

The software was developed in the Arduino programming environment. The freely available Arduino Integrated Development Environment (IDE) provides functions and libraries for communicating and programming the Teensy boards. Thus, our code can be easily adapted to the specific requirements of each experiment.

### 2.2. Experimental design

We conducted four experiments to validate the frequency resolution and end phase of the stimuli generated with the LED display. In the first two experiments, we measured the stimuli’s frequency resolution and end phase using a photodiode-based circuit. In the last two experiments, we validated the frequency resolution and end phase of the visual stimuli by recording the EEG of human subjects who were exposed to the stimuli.

#### 2.2.1. Experiments 1 and 2

For experiment 1, we generated sinusoidal stimuli with frequencies from 9.5 to 10.5 Hz, with increments of 0.1 Hz and end phase of the stimuli of 0°. For experiment 2, we generated sinusoidal stimuli with a fixed frequency of 10 Hz and end phases of 0°to 315°, with increments of 45°.

For experiments 1 and 2, we measured the entrainer LEDs light intensities with a transimpedance amplifier [23] (see Fig. 2). The amplifier uses a VTB8440 (Excelitas Technologies, MA, USA) visible light photodiode and an MCP6241 operational amplifier (Microchip Technology Incorporated, AZ, USA). We adjusted the gain of the amplifier to get an output voltage of 5 V when the LEDs were on. We recorded the amplifier output voltage with a Saleae Logic 8 (Saleae, CA, USA) with a sampling frequency of 25 MHz.

**Figure 2.**
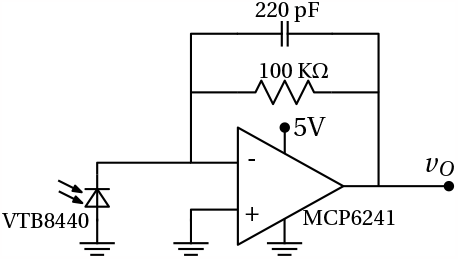
Light intensity measurement circuit. The LEDs light intensity is measured using a photodiode and a transimpedance amplifier. A logic analyzer measures the amplifier voltage.

For experiment 1, we measured the stimulation frequency by performing a spectral analysis of the recorded LEDs light intensities. We estimated the recording’s power spectral density (PSD) using Welch’s method with a Hamming window [24], using the MATLAB2021b implementation. The spectral analysis parameters were defined according to the desired physical frequency resolution Δ *f* given by

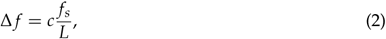

where *c, f*_*s*_, and *L* are the window-dependent constant (*c* = 2 for a Hamming window), sampling rate, and window length, respectively [25]. We chose Δ *f* = 0.05 Hz and 5 segments with 50% overlap, leading to recordings with a total duration of 60 seconds.

For experiment 2, we measured the end phase *ϕ*_END_ of the stimulation by decoding the last two pulses of the LEDs light intensity measurement. First, we computed the duty cycle of these last two pulses (see Fig. 5-(a)) as

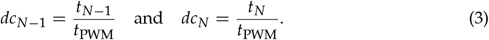

Solving for *ϕ* in Eq. (1) we get

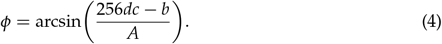

The sine function is not one-to-one, thus, the range of its inverse is limited to the [− 90^*°*^, 90^*°*^] interval. Since we are interested in angles in the range [0^*°*^, 360^*°*^], we used the sign of the duty cycle slope to compute the end phase:

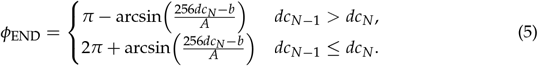

#### 2.2.2. Experiments 3 and 4

Experiments 3 and 4 were conducted to show the effect of visual stimulation generated by our LED display on four participants (see Fig. 3). Experiment 3 is related to the generation of sinusoidal-like stimulation at different frequencies, and experiment 4 is related to the variation of the phase at the stimulation offset.

**Figure 3.**
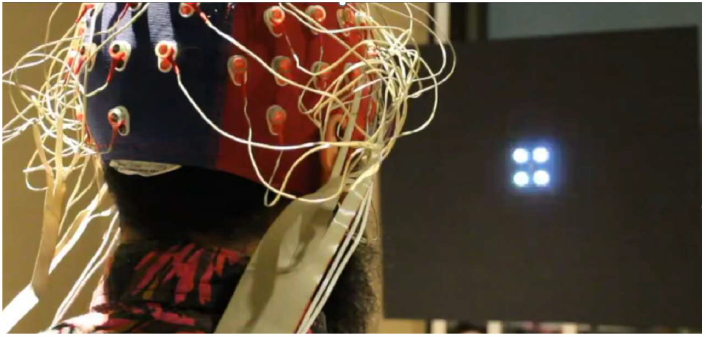
Image from the LED display during an SSVEP experiment.

The EEG recordings were conducted on four healthy participants following the experimental protocol described in [19]. Before joining the study, all participants provided their written consent by signing a consent form. The experimental procedure was approved by the Research and Ethics Committee of the Universidad de Valparaíso (evaluation statement code CEC170-18), and adhered to the national regulations for human subject research and the principles outlined in the Declaration of Helsinki.

The participants were seated within a dimly illuminated, acoustically insulated, and electromagnetically shielded EEG chamber. Eyes-open resting-state EEG data was recorded for 5 minutes, from 64 scalp locations, using an Active II Biosemi acquisition system. EEG signal was sampled at 8 kHz and processed offline using Brain Vision Analyzer 2.0 (Brain Products GmbH, Munich, Germany) using the steps recommended in [26]. Data was segmented into epochs of 5 s. The fast Fourier Transform (FFT) was calculated for each segment using a frequency resolution of 0.125 Hz, and the mean power spectrum was computed [27]. The frequency with the highest power spectrum value in the 8–14 Hz frequency band at the electrode Oz was determined to set the individual alpha frequency (IAF) [28,29] for each subject.

The IAF was the frequency selected for the visual stimulation and the generation of SSVEP. To this end, the led display was placed at 70 cm from the participant. The vertical position of the LED display was adjusted to align with the eye level of the participant, who was instructed to focus on the center of the LED display.

During stimulation, the light intensity of the four LEDs followed a synchronized sinusoidal pattern of waxing and waning, matching the IAF of each participant [19]. We adjusted the maximum intensity of the stimulation according to the participant’s comfort, maintaining a suprathreshold minimum intensity. Stimuli were presented in three blocks of 60 trials each. EEG signals were processed similarly to the description provided above for resting-state EEG. Recordings were analyzed in the frequency domain and the power of the SSVEP was defined as the mean power spectrum between trials.

Experiment 4 consisted of four experimental conditions in which the stimulus was set to finish at one of four possible phases of the sinusoid: 0°, 90°, 180°, and 270°as in [19]. Noteworthy, the stimulus onset did not vary among experimental conditions and always started at phase zero, i.e., mean intensity in the ascending ramp of the sinusoid.

## 3. Results

### 3.1. Experiments 1 and 2

Figure 4 shows the estimated power spectral densities of the recorded signals during experiment 1. We can observe a clear peak in the PSD at the corresponding stimulation frequencies. Table 1 shows the results for experiment 2, including the nominal end phase *ϕ*_END_ (for values 0°, 45°, …, 315°), the measured end phase 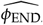, and the absolute end phase error error_*ϕ* END_, all in degrees. The table also shows the corresponding PWM values (obtained through Eq. (1)) for the nominal value PWM_END_, the measured value 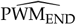value, and absolute error value error_PWM_. Fig. 5-(b) shows the end phase error value error_*ϕ* END_ (upper panel) and the PWM error value error_PWM_ (lower panel) for each end phase value. Note that there is a nonlinear relationship between error_*ϕ* END_ and error_PWM_ given by Eq.(5).

**Table 1.**
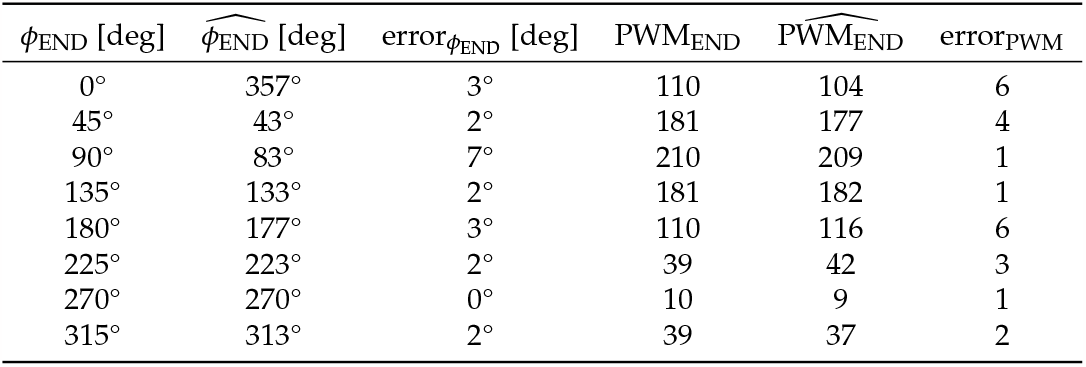
Measured end phases and corresponding PWM values.

| $\phi_{\text{END}}$ [deg] | $\widehat{\phi}_{\text{END}}$ [deg] | $\text{error}_{\phi_{\text{END}}}$ [deg] | $\text{PWM}_{\text{END}}$ | $\widehat{\text{PWM}}_{\text{END}}$ | $\text{error}_{\text{PWM}}$ |
| --- | --- | --- | --- | --- | --- |
| 0° | 357° | 3° | 110 | 104 | 6 |
| 45° | 43° | 2° | 181 | 177 | 4 |
| 90° | 83° | 7° | 210 | 209 | 1 |
| 135° | 133° | 2° | 181 | 182 | 1 |
| 180° | 177° | 3° | 110 | 116 | 6 |
| 225° | 223° | 2° | 39 | 42 | 3 |
| 270° | 270° | 0° | 10 | 9 | 1 |
| 315° | 313° | 2° | 39 | 37 | 2 |

**Figure 4.**
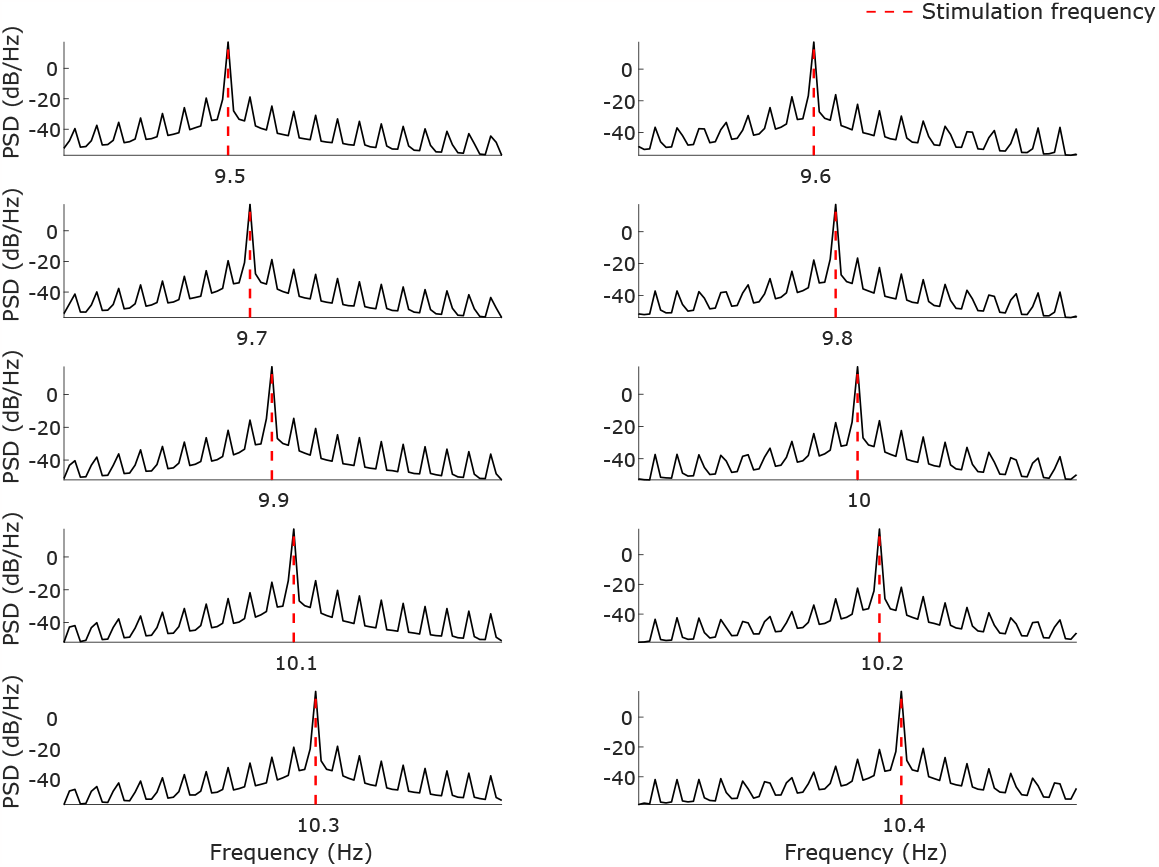
PSD estimation of the recorded LEDs light intensities during experiment 1. The PSDs were estimated using Welch’s method. The stimulation frequencies were set from 9.5 to 10.4 Hz with increments of 0.1 Hz.

**Figure 5.**
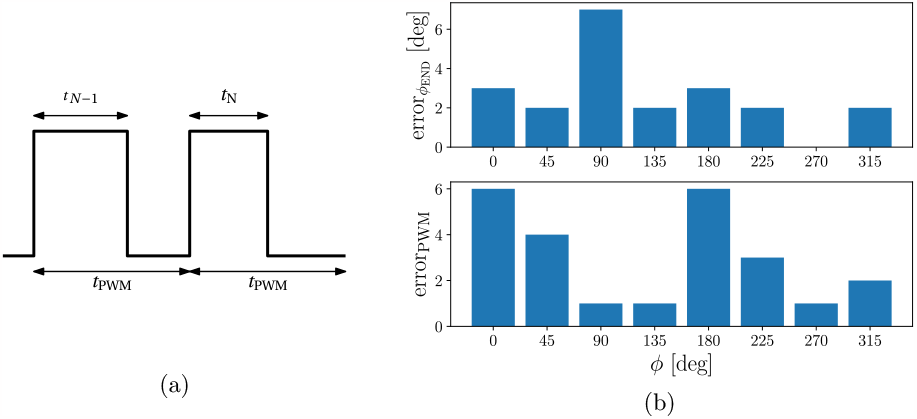
Phase error: (a) The phase is computed using the duty cycle of the last two PWM pulses 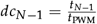 and 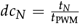, respectively. (b) Measured PWM error (error_PWM_) and phase error (error_*ϕ*_) for reference phases 0°, 45°, …, 315°.

### 3.2. Experiments 3 and 4

Our results showed that the stimulation generated the SSVEP at the expected frequency for all the study participants. Figure 6 shows the mean power spectral density in a pull of occipital electrodes (O1, O2, Oz, and POz) computed using the EEG recorded from four participants at rest (black lines) and during the visual stimulation (grey lines) according to the methodology of section 2.2.2.

**Figure 6.**
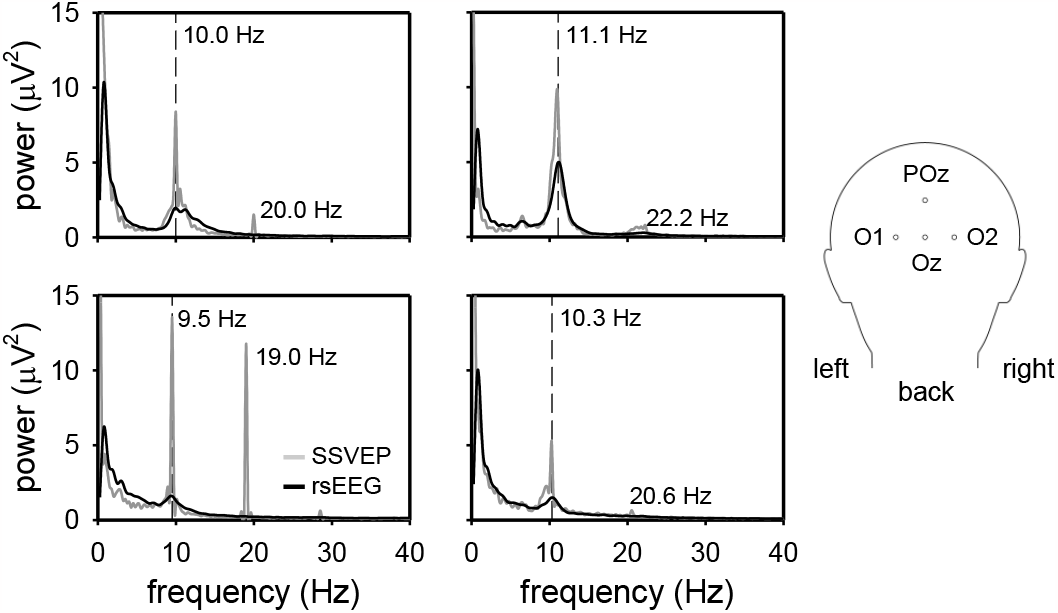
Mean of the power spectrum of four electrodes (O1,O2, POz and OZ) from four participants exposed to visual stimulation generated with the LED display. Inset at the right panel illustrates the position of the four electrodes in the conventional 10–20 EEG electrode placement scheme used in this study. Black and grey lines show the power spectrum of the resting-state EEG and the SSVEPs, respectively.

We could effectively generate SSVEP at the participant’s alpha frequency using the LED screen. In Figure 6 black lines) we can observe the IAF, i.e., the peak in the range of 8-14Hz for each participant. During stimulation, a visible increase in power at the IAF and its first harmonics can be observed (Figure 6 grey lines). This result is in accordance with the SSVEP literature [1,12,13].

Results obtained from EEG recordings using this display to control the phase of stimulus at the offset of stimulation can be found at [19]. In this work, the specific phase angles of the recorded EEG signal at this offset were computed. The study revealed that the EEG signal phases at the end of the stimulus were dispersed over a consistent range of values compared to the phase at the stimulus’s offset (see [19] for more details).

## 4. Discussion

In this article, we presented a high-resolution, affordable, and customized LED display for visual stimulation to generate SSVEPs. Our approach uses LEDs, the DDS and PWM techniques to generate a sinusoidal-like signal to control the light intensity and phase of the visual stimulation. The LED display here presented is an open-hardware and software device [30] as we provide the hardware schematics and all the display software to promote the use of this approach and the replication of the study. The choice of Teensy as the display board aligns with this goal, as it can be programmed in the Arduino environment [31,32], and is compatible with most operating systems [32]. This selection not only contributes to cost-effectiveness but also ensures excellent performance. Various initiatives for low-cost instrumentation have also opted for Teensy as their board controller of choice, as observed in [33–35].

### 4.1. Comparison with other available systems

Other proposals for displays with visual stimulation purposes have been made using LED technology. In [36], a stimulus for the fly visual system was designed to present apparent motion stimuli. This display technology was used to perform behavioral experiments with Drosophila. Furthermore, in the work of [37], a technique was developed using frequency-modulated visual stimuli for brain–computer interfaces, adapting the stimulation approach usually used in the auditory domain to evoke steady-state responses. In this approach, only 10 Hz was used as a modulated signal, and the SSVEPs recorded with EEG were compared to flicker stimulation in terms of perceptibility ratings.

In [38], a LED display was created using an Arduino microcontroller for visual stimulation. This study demonstrated low latencies (1.2-2.4 ms) and established a linear relationship between the PWM duty cycle and the measured normalized light output, making this technique suitable for neuroscience experiments involving cue light tasks. The authors validated their system by assessing the light output quality and switching times. Compared to the system proposed here, a disadvantage of the approach in [38] is the computer requirement during experiment execution and the absence of triggers to mark the LED stimulation’s starting point. However, it is essential to note that this system was not designed for the sinusoidal modulation of LEDs, as the display presented here.

The system proposed in [39] was mainly focused on ensuring low power consumption for long-lasting battery applications. In the case of [39], unlike the sinusoidal-like stimulation proposed in this work, they used flickered signals at a rate of 20 Hz (with a fixed duty cycle of 50%). This approach does not allow the control of the phase of the stimulation.

### 4.2. Applications within and beyond visual stimulation

Another important aspect of the proposed system is its versatility in terms of applications. This system is ready to use it in experiments to generate SSVEPs. The current system also provides an additional LED at the center of the square, composed of the four driver LEDs, which could present transient visual stimulation at any time during the periodic stimulation. It is also possible to adapt the system to work with tactile stimulation delivered by solenoid stimulators, such as the Dancer Design tactor [40], with a minor change in the driver circuit, to generate Steady State Somatosensory Evoked Potentials [41,42].

## 5. Conclusion

In this article an open-source LED display with a simple and consequently affordable controlling system is presented. Our system allows the definition of arbitrary stimulation frequencies with a 0.1 Hz resolution and offers control of the phase of the stimulus. Moreover, the LED display is also prepared to send triggers at the begining (onset) and the end (offset) of the stimulation ensuring synchronization between the recording system and the stimulation. This LED display system could be useful in several possible experimental paradigms. Given that the software and circuits are accessible to the entire research community, the system can be personalized and configure to meet the diverse needs of different research hypothesis, making it a versatile solution for various scientific projects.

## Author Contributions

Conceptualization, M.O, W.E. and A.W.; methodology, software and hardware, M.O, Y.P.C. and A.W; validation, M.O and Y.P.C.; data curation, M.O; writing—original draft preparation, M.O, A.W., and Y.P.C. All authors contributed to manuscript revision and agreed to the published version of the manuscript.

## Institutional Review Board Statement

The studies involving human participants were reviewed and approved by Research and Ethics Committee of the Universidad de Valparaíso (assessment statement code CEC170-18, June 19th, 2018).

## Informed Consent Statement

The participants provided their written informed consent to participate in this study.

## Data Availability Statement

https://doi.org/10.5281/zenodo.10223153

## Abbreviations

The following abbreviations are used in this manuscript:

LED: Light-Emitting Diode
EEG: Electroencephalogram
SSVEP: Steady State Visual Evoked Potential
PWM: Pulse Width Modulation
DDS: Direct Digital Synthesis
PSD: Power Spectral Density
IDE: Integrated Development Environment
IAF: Individual Alpha Frequency

## Disclaimer/Publisher’s Note

The statements, opinions and data contained in all publications are solely those of the individual author(s) and contributor(s) and not of MDPI and/or the editor(s). MDPI and/or the editor(s) disclaim responsibility for any injury to people or property resulting from any ideas, methods, instructions or products referred to in the content.s

